# PQN-59 antagonizes microRNA-mediated repression and functions in stress granule formation during *C. elegans* development

**DOI:** 10.1101/2021.05.14.444139

**Authors:** Colleen Carlston, Robin Weinmann, Natalia Stec, Simona Abbatemarco, Francoise Schwager, Jing Wang, Huiwu Ouyang, Monica Gotta, Christopher M. Hammell

**Author notes:** Boston Children’s Hospital, Boston, MA 02115. DKFZ & BioQuant Center, Heidelberg, 69120 Germany. Correspondence (C.M.H.).

## Abstract

microRNAs (miRNAs) are potent regulators of gene expression that function in a variety of developmental and physiological processes by dampening the expression of their target genes at a post-transcriptional level. In many gene regulatory networks (GRNs), miRNAs function in a switch-like manner whereby their expression and activity elicit a transition from one stable pattern of gene expression to a distinct, equally stable pattern required to define a nascent cell fate. While the importance of miRNAs that function in this capacity are clear, we have less of an understanding of the cellular factors and mechanisms that ensure the robustness of this form of regulatory bistability. In a screen to identify suppressors of temporal patterning phenotypes that result from ineffective miRNA-mediated target repression during *C. elegans* development, we identified *pqn-59,* an ortholog of human UBAP2L, as a novel factor that antagonizes the activities of multiple heterochronic miRNAs. Specifically, we find that depletion of *pqn-59* can restore normal development in animals with reduced miRNA activity. Importantly, inactivation of *pqn-59* is not sufficient to bypass the requirement of these regulatory RNAs within the heterochronic GRN. The *pqn-59* gene encodes an abundant, cytoplasmically localized and unstructured protein that harbors three essential “prion-like” domains. These domains exhibit LLPS properties *in vitro* and normally function to limit PQN-59 diffusion in the cytoplasm *in vivo*. Like human UBAP2L, PQN-59’s localization becomes highly dynamic during stress conditions where it re-distributes to cytoplasmic stress granules and is important for their formation. Proteomic analysis of PQN-59 complexes from embryonic extracts indicates that PQN-59 and human UBAP2L interact with orthologous cellular components involved in RNA metabolism and promoting protein translation and that PQN-59 additionally interacts with proteins involved in transcription and intracellular transport. Finally, we demonstrate that *pqn-59* depletion results in the stabilization of several mature miRNAs (including those involved in temporal patterning) without altering steady-state pre-miRNAs levels indicating that PQN-59 may ensure the bistability of some GRNs that require miRNA functions by promoting miRNA turnover and, like UBAP2L, enhancing protein translation.

**AUTHOR SUMMARY:** Bistability plays a central role in many gene regulatory networks (GRNs) that control developmental processes where distinct and mutually exclusive cell fates are generated in a defined order. While genetic analysis has identified a number of gene types that promote these transitions, we know little regarding the mechanisms and players that ensure these decisions are robust. and in many cases, irreversible. We leveraged the robust genetics and phenotypes associated with temporal patterning mutants of *C. elegans* to identify genes whose depletion would restore normal regulation in animals that express miRNA alleles that do not sufficiently down-regulate their targets. These efforts identified *pqn-59*, the *C. elegans* ortholog of the human UBAP2L gene. Like UBAP2L, PQN-59 likely forms a hub for a number of RNA/RNA-binding protein mediated processes in cells including translational activation and in the formation of stress granules in adverse environmental conditions. Finally, we also demonstrate that *pqn-59* depletion stabilizes mature miRNA levels further connecting this new family of RNA-binding proteins to translation and miRNA-mediated gene regulation.

## INTRODUCTION

Cell fate specification during animal development is tightly controlled to yield highly reproducible outcomes and avoid extreme variation. While individual cell fates can be described by quantifying the expression levels of RNAs that are expressed at any given time, the stable patterns of gene expression for a distinct cell type are often governed by the presence and levels of a relatively limited number of master regulator genes that function at the top of a gene regulation hierarchy. These master regulatory genes typically function in a concentration-dependent manner whereby expression levels above a critical threshold are sufficient to program downstream patterns of transcription in a dominant way. Changes in cell fate specification can occur in a switch-like manner when intrinsic or extrinsic changes lead to reduced expression or activity of a master regulator to levels below a discrete, critical threshold [1]. What common design features of GRNs set these thresholds and what cellular components ensure that the outcomes of sharp changes in the levels of master regulator genes are translated into distinct cell fates have been understudied.

The remarkable precision of *C. elegans* post postembryonic cell fate specification relies on GRNs that exhibit switch-like behavior to ensure normal temporal development. Larval development proceeds through four stages (separated by molts) where the timing of individual cell divisions and the patterning of temporal cell fates is invariant in wild-type animals [2]. Heterochronic genes, encoding transcription factors (TFs) and RNA-binding proteins, organize the sequence of temporal development through the control of stage-specific patterns of gene expression [3]. Importantly, these factors function in dosage-sensitive manners and exhibit sharp temporal gradients of expression (usually from high expression to low expression) that change during inter stage molts. miRNAs play a central role in promoting these sharp transitions from one stage to the next by curtailing the expression of discrete protein-coding genes of the heterochronic pathway that define individual temporal cell fates. This process occurs in at least three separate phases of development (L1-to-L2, L2-to-L3, and L4-to-adulthood) and is mediated by distinct miRNAs that regulate the translational output of independent target mRNAs [4]. Importantly, heterochronic miRNAs also function in dosage-sensitive manners whereby alterations in activity or the timing of their expression lead to the perdurance of target gene expression; resulting in the inappropriate reiteration of earlier patterns of cell division and cell fate specification at subsequent molts [4].

In this study, we aimed to identify candidate genes that modulate the ability of heterochronic miRNAs to elicit switch-like reprograming of temporal cell fates. Specifically, we sought to identify genes whose inactivation would enable hypomorphic alleles of miRNA genes that alone are incapable of dampening the expression of their mRNA targets below a required threshold to function normally. These efforts identified *pqn-59*, encoding a previously uncharacterized, cytoplasmically localized “prion-like” domain-containing protein, as a gene product whose inactivation suppresses loss-of-function phenotypes associated with ineffective miRNA-mediated repression. Consistent with *pqn-59* functioning as a modulator of heterochronic miRNA function in temporal cell fate switching, *pqn-59* cannot bypass the requirement for these regulatory RNAs in the heterochronic GRN. We demonstrate that the PQN-59 protein exhibits liquid-liquid phase separation (LLPS) properties *in vivo* and *in vitro* and that PQN-59 is important for the assembly of RNA/RNA-binding protein assemblies called stress granules during heat shock; a feature shared with its human ortholog UBAP2L. A comparison of proteins associated with PQN-59 indicate that PQN-59, like UBAP2L, is likely highly integrated with proteins involved in mRNA metabolism and translational activation. Finally, we show that *pqn-59* depletion may suppress temporal patterning defects in miRNA mutants by increasing the steady state levels of miRNAs in addition to its potential roles in translational activation. In this capacity, normal PQN-59 activity functions to set and maintain gene expression levels within the range for executing bistable switches in gene expression required for normal development.

## RESULTS

### *pqn-59* depletion suppresses the heterochronic phenotypes of *lin-4* loss-of-function mutants

In order to identify additional gene products that modulate heterochronic miRNA-dependent developmental events, we performed a genome-wide suppressor screen to identify genes whose inactivation via RNAi could suppress the reiterative heterochronic phenotypes of a hypomorphic allele of the *lin-4* miRNA gene. The *ma161* allele of *lin-4* is defined by a single nucleotide substitution in the mature *lin-4* miRNA that reduces its ability to down-regulate its target mRNA, *lin-14,* which is normally turned off by the second larval stage [5]. As a consequence of these molecular defects, *lin-4(ma161)* animals continually express LIN-14 throughout larval development and reiterate L1 specific patterns of cell differentiation for each of the somatic blast cells at subsequent molts. Importantly, while the *lin-4(ma161)* allele generates a small regulatory RNA, its developmental phenotypes are indistinguishable from those completely lacking the *lin-4* gene [5, 6]. As a consequence of these temporal patterning defects, *lin-4(ma161)* animals lack vulval structures required for normal egg laying and also fail to induce the expression of an adult-specific *col-19::GFP* transcriptional reporter after the fourth larval molt (Figure 1A and B). To identify suppressors, we exposed *lin-4(ma161)* animals harboring the *col-19::GFP* reporter to individual clones of a genomic scale RNAi library and identified dsRNAs that could restore normal *col-19::GFP* expression during adulthood. One of these clones generated dsRNA against *pqn-59*, a highly conserved and uncharacterized *C. elegans* gene, that robustly suppressed the reiterative heterochronic phenotypes of lin-4(ma161) mutants to a similar level as other previously described suppressors (including *lin-14, lin-28*, and *lin-42*)(Table 1).. Examination of adult *lin-4(ma161*) animals that had been exposed to *pqn-59* dsRNAs exhibited normal temporal seam cell developmental division patterns and were now able to generate alae production on adult cuticles, indicative of normal seam cell temporal patterning (Figure 1C). Furthermore, in contrast to control RNAi animals, *pqn-59* RNAi also suppressed the vulvaless phenotypes of *lin-4(ma161)* animals, enabling these animals to lay eggs (Figure 1D). Surprisingly, depletion of *pqn-59* activity in wild-type animals did not induce precocious deposition of adult-specific alae at the L4 molt which distinguishes it from other previously characterized *lin-4* suppressors) (Table 1). Furthermore, *pqn-59* depletion in wild-type backgrounds only induced a mild, early expression of the *col-19::GFP* reporter in hypodermal cells found in the head and tail regions (H0, H1, and T cells) of approximately 17% of late L4-staged animals (L4.5 or later [7]).

**Figure 1.**
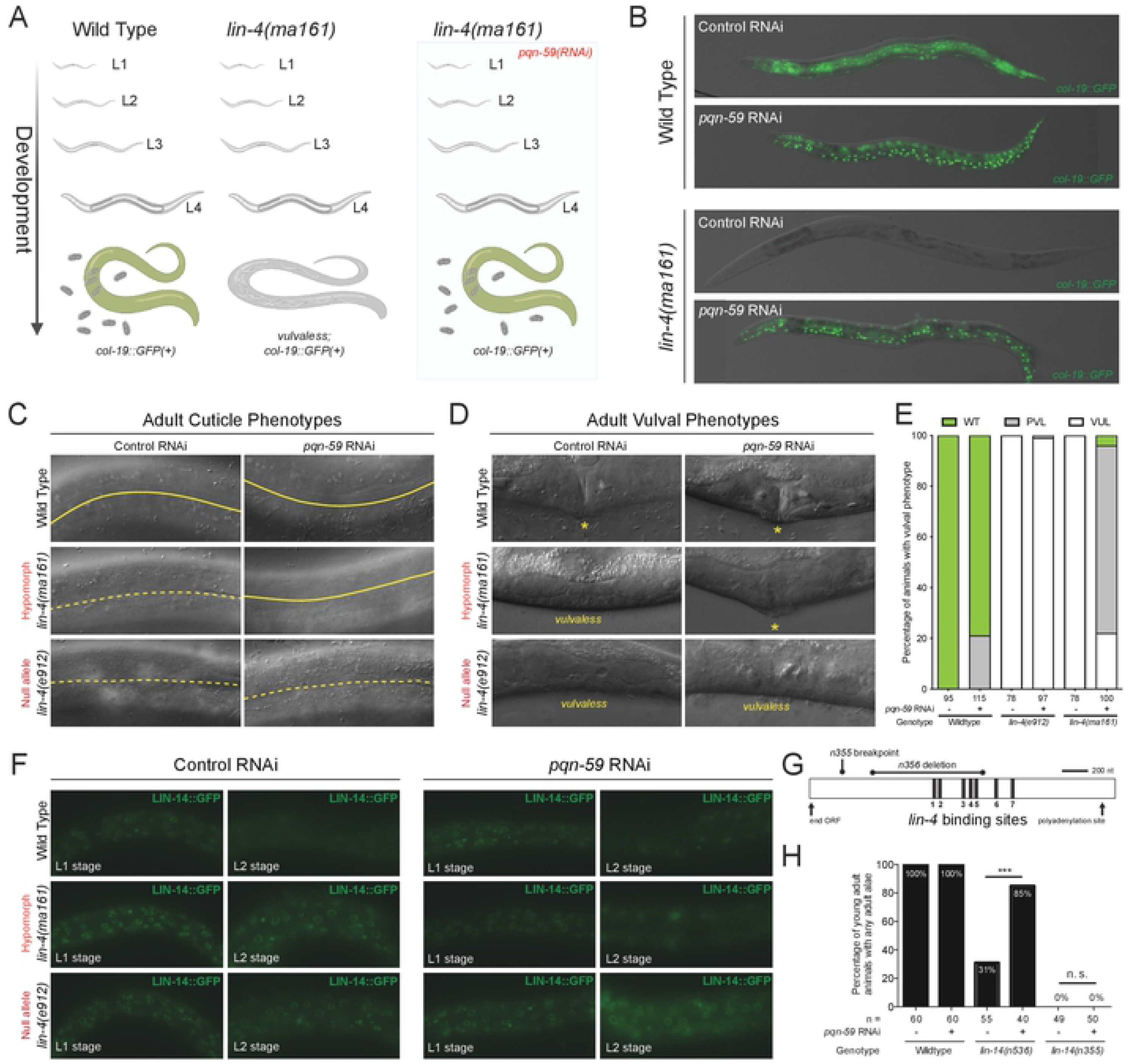
*pqn-59(RNAi)* suppresses reiterative heterochronic phenotypes associated with reduced *lin-4*-mediated repression of *lin-14.* **(A)** Genetic screen used to identify suppressors of *lin-4(ma161)* heterochronic phenotypes. Wild-type animals express the adult-specific *col-19::GFP* transcriptional reporter at the end of the L4 molt and throughout adulthood. In contrast, *lin-4(ma161)* animals fail to express *col-19::GFP* and are unable to lay eggs due to the inappropriate reiteration of L1-specific developmental programs. Depletion of *pqn-59* by RNAi suppresses both *lin-4(ma161)* phenotypes. **(B)** Pictographs of adult wild-type and *lin-4(ma161)* animals exposed to control or *pqn-59* dsRNAs. See Table 1 for details. **(C and D)** Pictographs depicting the comparison of adult-specific alae and vulval structures of wild-type, *lin-4(ma161),* and *lin-4(e912)* animals exposed to control or *pqn-59* dsRNAs. In panel C, solid line indicates continuous adult alae while a dashed line depicts the regions of the cuticle that lack these cuticular structures. In panel D, asterisks (*) indicate the location of a normal vulval structures in RNAi treated animals at the L4 to adult molt. **(E)** Quantification of the wild-type, protruding vulva (*pvl*), and vulvaless (*vul*) phenotypes of the indicated animals treated with control or *pqn-59* dsRNAs. **(F)** Images depicting the temporal expression patterns of an endogenous, GFP-tagged allele of the protein product of the major lin-4 target, LIN-14. **(G)** Diagram indicating the genetic lesions that alter the *lin-14* 3’ UTR regulatory sequences. Black bars indicate the approximate locations of the complementary *lin-4* binding sites in the *lin-14* 3’UTR. **(H)** Quantification of the levels of suppression mediated by depleting *pqn-59* during the development of animals the express one of two gain-of-function alleles of *lin-14*. Animals were scored positive if any visible alae structures were present on the adult-stage cuticle.

**Table 1.** Measurement of RNAi-mediated suppression of temporal patterning phenotypes of various heterochronic mutants. This table contains the quantification of alae and *col-19::GFP* expression phenotypes of various heterochronic mutants with and without *pqn-59*(RNAi).

| # | Strain | Genotype<sup>a</sup> | Experimental Condition<sup>b</sup> | % Animals L4 alae<sup>c</sup> | n = | % Animals Adult alae<sup>c</sup> | n = | % Animals expressing adult *col-19::GFP* | n = |
| --- | --- | --- | --- | --- | --- | --- | --- | --- | --- |
| 1 | VT1367 | Wild Type | control | 0 | 80 | 100 | 100 | 100 | 100 |
|  |  |  | *pqn-59* RNAi | 0 | 110 | 100 | 100 | 100 | 100 |
|  |  |  | *lin-42* RNAi | 19 | 100 | 100 | 100 | 100 | 100 |
|  |  |  | *lin-14* RNAi | 83 | 100 | 100 | 100 | 100 | 100 |
|  |  |  | *lin-28* RNAi | 84 | 100 | 100 | 100 | 100 | 100 |
| 2 | HML2 | *lin-4(ma161)* | control | 0 | 100 | 0 | 80 | 0 | 100 |
|  |  |  | *pqn-59* RNAi | 0 | 100 | 56 | 100 | 62 | 100 |
|  |  |  | *lin-42* RNAi | 0 | 100 | 46 | 100 | 77 | 100 |
|  |  |  | *lin-14* RNAi | 4 | 100 | 81 | 100 | 73 | 100 |
|  |  |  | *lin-28* RNAi | 16 | 100 | 89 | 100 | 89 | 100 |
| 3 | HML6 | *lin-4(e912)* | control | - | - | 0 | 32 | 0 | 30 |
|  |  |  | *pqn-59* RNAi | - | - | 0 | 30 | 5 | 30 |
| 4 | HML11 | *let-7(n2853)* | control | - | - | - | - | 35<sup>d</sup> | 100 |
|  |  |  | *pqn-59* RNAi | - | - | - | - | 60<sup>d</sup> | 100 |
|  |  |  | *lin-42* RNAi | - | - | - | - | 61<sup>d</sup> | 100 |
|  |  |  | *lin-14* RNAi | - | - | - | - | 58<sup>d</sup> | 100 |
|  |  |  | *lin-28* RNAi | - | - | - | - | 84<sup>d</sup> | 100 |
| 5 | VT2077 | *lin-31(n1053)* | control | - | - | - | - | 100 | 100 |
|  |  |  | *pqn-59* RNAi | - | - | - | - | 100 | 100 |
|  |  |  | *lin-42* RNAi | - | - | - | - | 100 | 100 |
|  |  |  | *lin-14* RNAi | - | - | - | - | 100 | 100 |
|  |  |  | *lin-28* RNAi | - | - | - | - | 100 | 100 |
| 6 | VT2079 | *lin-31(n1053); alg-1(ma192)* | control | - | - | - | - | 7 | 100 |
|  |  |  | *pqn-59* RNAi | - | - | - | - | 62 | 100 |
|  |  |  | *lin-42* RNAi | - | - | - | - | 26 | 100 |
|  |  |  | *lin-14* RNAi | - | - | - | - | 66 | 100 |
|  |  |  | *lin-28* RNAi | - | - | - | - | 64 | 100 |

<sup>a</sup>Animals contain *mal-105* containing a integrated *col-19::GFP* transgenic array
<sup>b</sup>P<sub>0</sub> animals were exposed to bacteria expressing indicated dsRNAs and adult F1 animals were scored for the indicated phenotype
<sup>c</sup>Presence and quality of cuticular alae structures were assayed by Normarski DIC optics. Only one side of each animals was scored.
<sup>d</sup>*let-7(n2853)* animals exhibit some *col-19::GFP* expression in lateral seam cells and were scored positive if there was any expression in any hypodermal cells.
When animals were treated with either *pqn-59*, *lin-42*, *lin-14*, or *lin-28* dsRNAs, there was a clear difference in the expressivity of the suppressed phenotype with most positive scoring animals expressing *col-19::GFP* in seam and hyp7 cells.

### Suppression of *lin-4* loss of function phenotypes by *pqn-59(RNAi)* requires miRNA expression and *lin-4* miRNA target sites in the *lin-14* mRNA

Because the *lin-4(ma161)* allele generates some *lin-4* miRNA, we asked whether depleting *pqn-59* via RNAi could completely bypass the requirement for *lin-4* during post-embryonic development. To accomplish this, we repeated RNAi-based experiments in animals lacking the *lin-4* gene (e.g., *lin-4(e912)*). These experiments revealed that *pqn-59* depletion had little effect on the *col-19::GFP* expression in *lin-4(0)* mutants (Table 1). Furthermore, adult-specific alae formation and suppression of the vulvaless phenotypes were not restored in this genetic context indicating that *pqn-59(RNAi)* cannot bypass the normal requirement for *lin-4* in temporal patterning (Figure 1C and Table 1). We then aimed to determine how *pqn-59(RNAi)* modulates the temporal expression of LIN-14 protein, encoded by the major *lin-4* target mRNA [8]. We therefore examined the expression of a functional, CRISPR-tagged *lin-14::GFP* allele in wild-type, *lin-4(ma161)* and *lin-4(e912)* genetic backgrounds in experimental conditions where *pqn-59* expression is altered. In wild-type animals, LIN-14::GFP expression begins during embryogenesis and is post-transcriptionally down-regulated at the L1 to L2 molt by *lin-4* miRNAs [8, 9]. In contrast, LIN-14::GFP expression perdures throughout development in both *lin-4(ma161)* and *lin-4(e912)* animals (Figure 1F). When *lin-4(ma161)* animals were treated with *pqn-59* dsRNAs, LIN-14::GFP expression was curtailed by the early L2 stage with similar kinetics as observed in wild-type animals exposed to either control or *pqn-59* dsRNAs (Figure 1F). In contrast, LIN-14::GFP expression is continuously maintained in *lin-4(e912)* animals in these conditions indicating that *pqn-59* depletion does not bypass the requirement for this miRNA in development (Figure 1F). We did note that LIN-14 expression in embryos and in L1-staged animals was slightly lowered in *pqn-59*(RNAi) conditions.

To further this analysis, we examined alleles of *lin-14* that generate mRNAs that harbor deletions in the 3’UTR of the *lin-14* transcript and have a reduced ability to be targeted by the *lin-4* miRNA. Specifically, we examined whether *pqn-59(RNAi)* could suppress the reiterative, gain-of-function phenotypes of *lin-14(n536)* and *lin-14(n355)* which harbor deletions of the *lin-14* 3’UTR that result in the deletion of five of seven or all of the *lin-4* miRNA binding sites in the *lin-14* 3’UTR (Figure 1G) [8]. Each of these mutants exhibit *lin-4*-like phenotypes (with different penetrance) due to the reduced ability of the *lin-14* mRNAs produced from these loci to be down-regulated the lin-4 miRNA [5]. We found that *pqn-59(RNAi)* was able to restore adult-specific alae formation in *lin-14(n536)* mutant animals that retain two of the seven *lin-4* binding sites in the *lin-14* 3’ UTR but not a *lin-14* truncation allele, *lin-14(n355),* that lacks all of the *lin-4* binding sites (Figure 1H). These experiments suggest that the suppression of *lin-4* reiterative phenotypes mediated by reducing *pqn-59* expression requires the ability of *lin-4* to both be produced and also able to physically interact with its target mRNA via complementary binding sites.

### *pqn-59* depletion can suppress other heterochronic phenotypes associated with reduced miRNA activity

We next aimed to determine if the genetic interactions between *pqn-59* and *lin-4* hypomorphic alleles are specific to developmental events that occur between the L1 to L2 stages of temporal development or whether *pqn-59* plays a more general role in antagonizing the activities of additional miRNAs that function later in the heterochronic pathway. We depleted *pqn-59* expression in two strains that exhibit distinct, reiterative alterations in the post-embryonic cell lineage due to a reduction in the activity of genetically separate sets of temporally regulated miRNAs. Animals containing the *let-7(n2956)* allele of the *let-7* gene fail to properly down-regulate the expression of LIN-41 during late larval development and as a consequence, animals reiterate the L4 patterns of hypodermal cell divisions during adulthood [10, 11]. A single amino-acid substitution (S895F) in one of the two *C. elegans <u>mi</u>*cro*<u>R</u>*NA-induced *<u>s</u>*ilencing *<u>c</u>*omplex (miRISC) argonaute proteins, ALG-1, reduces the efficacy of the three *let-7*-family miRNAs (mir-48, miR-241, miR-84) that are required to down-regulate the expression of *hbl-1,* a transcription factor that is essential for L2-specific cell fates [12–14]. This regulatory defect leads to an inappropriate reiteration of L2-stage, proliferative seam cell divisions during the L3 stage [6, 15]. Both *let-7(n2853)* and *alg-1(ma192)* animals fail to express *col-19::GFP* after the L4-to-adult molt (Figure 2 and Table 1). Depletion of *pqn-59* suppresses these phenotypes indicting that *pqn-59* functions at multiple stages of post-embryonic development to control lateral seam cell fate specification (Figure 2A and B). Examination of the adult cuticles and the numbers of lateral seam of *alg-1(ma192)*; *pqn-59*(RNAi) animals indicated that reducing *pqn-59* activity during development can 1) restore the ability of these mutant animals to generate adult-specific alae and 2) limit the inappropriate reiteration of L2-stage seam cell division programs that leads to excessive proliferation of skin stem cells (Figure 2C).

**Figure 2.**
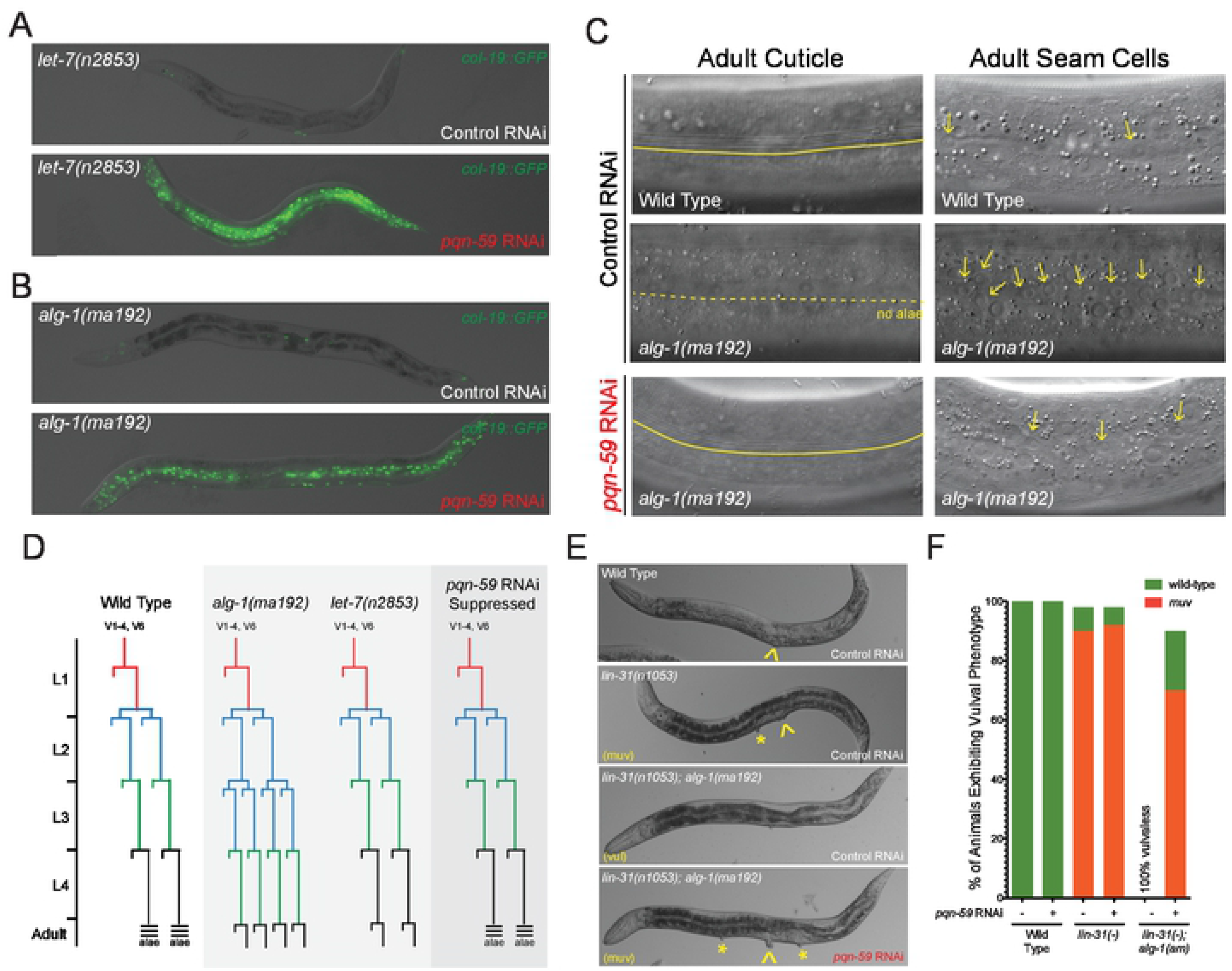
*pqn-59(RNAi)* suppresses additional miRNA-dependent reiterative heterochronic phenotypes. **(A and B)** Depletion of *pqn-59* via RNAi suppresses the *col-19::GFP* phenotypes of *let-7* hypomorphic alleles *(let-7(n2853))* and antimorphic alleles of genes that encode core components of miRISC (e.g., ALG-1, *alg-1(ma192))*. See Table 1 for details on expressivity and penetrance of suppression. **(C)** *alg-1(ma192)* mutants reiterate the L2-stage or larval development one additional time and as a consequence, over-proliferate lateral seam cells and also fail to form alae structures on the adult cuticle [6]. These alg-1(ma192)-dependent cell lineage phenotypes and defects in generating adult-specific alae are suppressed by depleting PQN-59 expression using dsRNAs against the *pqn-59* gene. Continuous yellow lines indicate regions of the cuticle where adult alae are present. Dashed lines indicate regions of the cuticle where alae are abnormally absent after the L4 molt. For images depicting the hypodermal cells of adult animals of indicated genotypes, arrows indicate the location of lateral seam cells. **(D)** Proposed seam cell linages of wild-type, let-7(n2853) and *alg-1(ma192)* animals after treatment with control or pqn-59 dsRNAs. **(E and F)** *alg-1(ma192)* animals exhibit a 100% penetrant synthetic vulvaless phenotype when combined with *lin-31* loss-of-function alleles, *lin-31(n1053)* [6, 15]. Depletion of *pqn-59* in *lin-31(n1053); alg-1(ma192)* double mutants completely suppresses these phenotypes to induce ectopic vulva in a *lin-31*-dependent manner.

*alg-1(ma192)* mutants also exhibit a synthetic phenotype with loss-of function mutations in the *C. elegans* HNF-3/fork head ortholog, *lin-31,* during vulval development. LIN-31 functions downstream of Ras/LET-60 signaling in vulval cell fate specification and loss-of-function alleles of *lin-31* (e.g., *lin-31(n1052)*) result in ectopic vulval induction of additional vulval precursor cells and a multi-vulval phenotype (Figure 2E and F) [16]. While *alg-1(ma192)* mutants generate a single vulval structure that bursts with high penetrance at the L4-to-adult molt, *lin-31(n1053); alg-1(ma192)* mutants exhibit a fully penetrant, synthetic vulvaless phenotype presumably due to alterations in the timing of vulval cell fate specification (Figure 2) [6, 15]. We found that, in addition to suppressing the reiterative seam cell phenotypes of *alg-1(ma192)* animals, *pqn-59* depletion almost completely suppressed the vulvaless phenotypes of *lin-31(n1053); alg-1(ma192)* double mutants (Figure 2E and F). This indicates that *pqn-59* also functions in the cell fate specification of vulval precursor cells.

### *pqn-59* encodes an essential protein that is localized in the cytoplasm throughout development

The *pqn-59* gene is located on the left arm of chromosome I and encodes a predicted 714 amino acid protein with a N-terminal *<u>ub</u>*iquitin-*<u>a</u>*ssociated (UBA) domain implicated structurally in various protein-protein interactions and in binding both ubiquitin and polyubiquitin chains (Figure 3A) [17]. Additional analysis of PQN-59 protein structure indicates that it also harbors two separate additional unstructured regions. Immediately after the N-terminal UBA-like domain, PQN-59 harbors a stretch of arginine/glycine-rich sequences (RGG/RG) that are typically associated with RNA binding proteins (Figure 3B) [18]. The C-terminal portion of PQN-59 harbors three “prion-like” domains composed of amino acid stretches disproportionately enriched for glutamine and asparagine residues (*<u>p</u>*rion-like, *<u>Q</u>* and *<u>N</u>*, *pqn-*) [19] (Figure 3B). A comparison of PQN-59 sequence to the proteomes of other model systems indicate that PQN-59 is evolutionarily related to a single *Drosophila* protein called Lingerer (Lig) and two human proteins named UBAP2 and UBAP2L (Figure 4C). Lig functions as a growth suppressor that associates with several RNA-binding proteins implicated in the regulation of protein translation (including Rasputin(Rin)/G3BP1 a RasGAP SH3 binding protein (RasGAP-BP), Caprin(Capr), and FMR1 an ortholog of the Fragile X mental retardation protein 1) [20]. Human UBAP2 and UBAP2L proteins have been demonstrated to promote translation [21] and are implicated in the formation of stress granules, liquid-liquid phase separating RNA-dependent condensates that form in the cytoplasm in a variety of averse cellular conditions [22–24].

**Figure 3.**
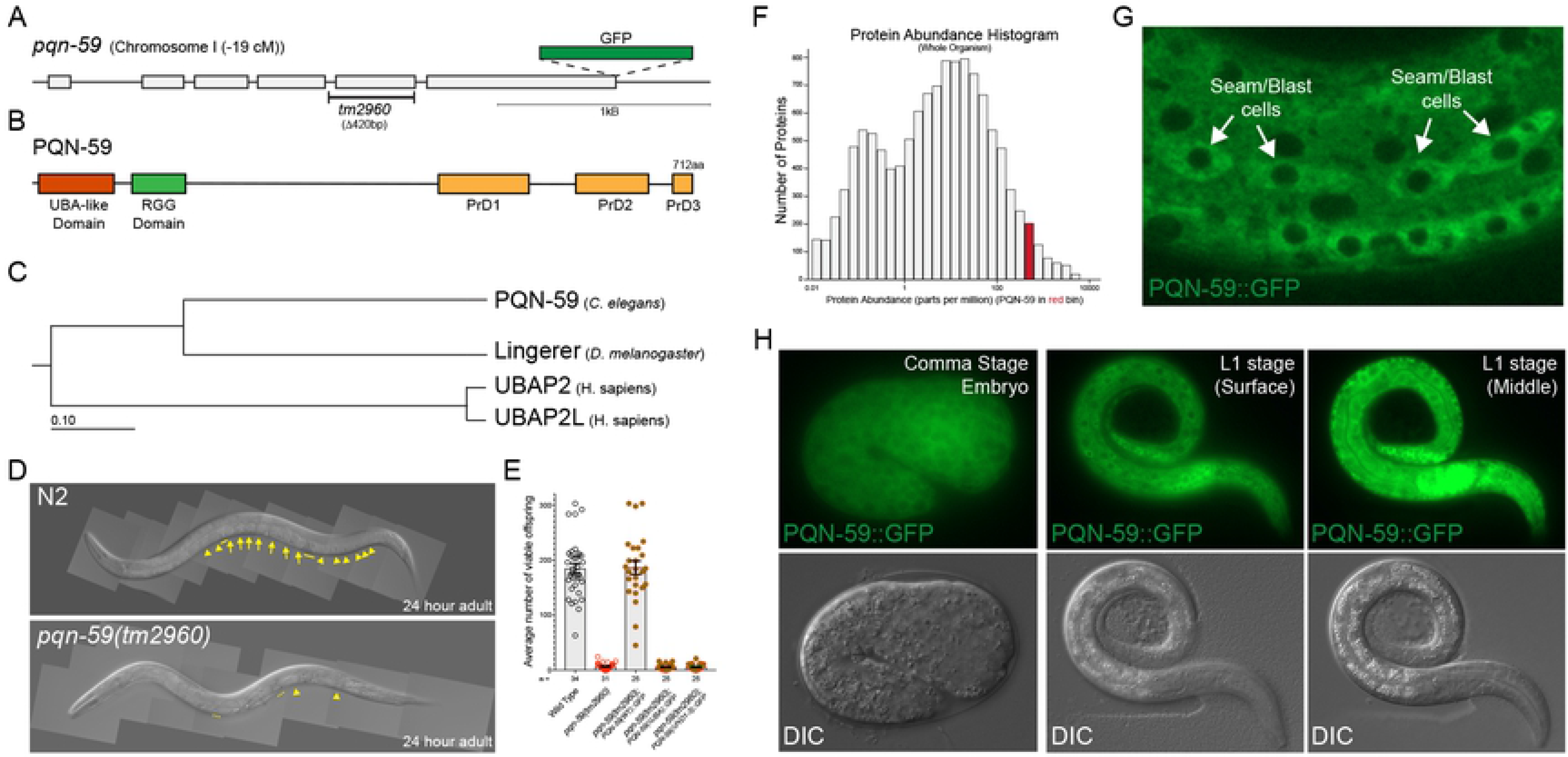
The *pqn-59* gene encodes an abundant, ubiquitously-expressed protein that is essential for normal fecundity. **(A and B)** Diagrams of the *pqn-59* gene and protein product. In panel A, the location of the *tm2960* deletion and the site of CRISPR-mediated GFP insertion that generates a functionally tagged allele of *pqn-59*. Panel B indicates the three discernable protein domains of PQN-59. **(C)** Evolutionary relationship between PQN-59 and orthologs in other systems. **(D)** Images of 24hr, adult wild-type and *pqn-59(tm2960)* animals demonstrating the differences in fertility. Arrowheads indicate developing oocytes and arrows indicate fertilized embryos. Dashed lines indicate regions of the germline where sperm are present. **(E)** Graph depicting the range of offspring with wild-type, *pqn-59(tm2960)* and *pqn-59(tm2960)* animals that express a functional, full-length PQN-59::GFP transgene (including upstream and downstream regulatory regions) targeted to chromosome II or variants of this transgene that lack the UBA or the three C-terminal prion-like domains. **(F)** Proteomic data from Version 4.0 of PaxDb: Protein abundance data indicating that PQN-59 is a very abundant protein. **(G)** Localization of the endogenously GFP-tagged allele of PQN-59 in lateral seam cells demonstrating that PQN-59 is a cytoplasmically localized. **(H)** Images of the same PQN-59::GFP transgene demonstrating that PQN-59 is expressed in all cell types throughout embryonic and post-embryonic development.

**Figure 4.**
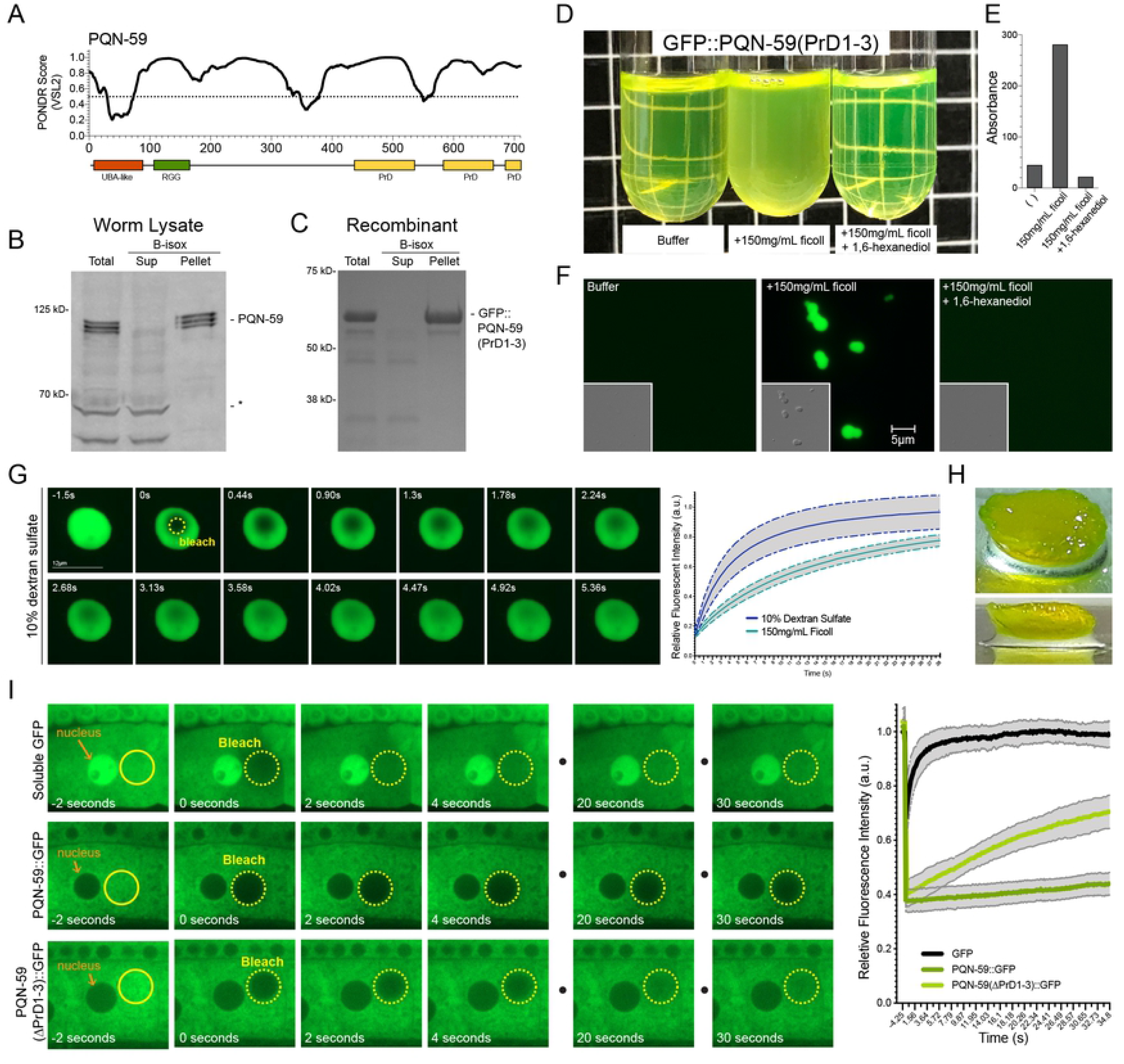
The “prion-like” domains of PQN-59 exhibit unique biopchysical properties *in vitro* and *in vivo*. **(A)** A majority of the PQN-59 protein is predicted to be unfolded using the Predictors of Natural Disorderd Regions (PONDR VSL2) algorithm [67]. **(B and C)** Endogenous, full-length PQN-59 in whole worm lysates and a GFP-tagged, C-terminal fragment of PQN-59 containing the three “prion-like” domains (PrDs) can be differentially co-precipitated with biotinylated-isoxazole (b-isox). **(D-F)** Solutions of purified GFP-PQN-59(PrD1-3) exhibit properties of LLPS when crowding reagents are added. Phase separation of GFP-PQN-59(PrD1-3) and increase absorbance phase separated samples are reversed upon addition of the aliphatic alcohol 1,6-hexandiol. **(G)** Phase separated droplets of GFP-PQN-59(PrD1-3) rapidly recover when photobleached. Graph depicts the rate of fluorescence recovery for GFP-PQN-59(PrD1-3) droplets that were formed in either dextran sulfate or ficoll concentrating agents. Graphs depict the average recovery rate of the GFP-PQN-59(PrD1-3) droplets in indicated concentrating agent (n = or greater than 10). Grey regions indicate standard error of measurement (SEM). **(H)** Prolonged concentration of GFP-PQN-59(PrD1-3) droplets results in the formation of a GFP-PQN-59(PrD1-3) hydrogel like material. **(I)** *In vivo*, full-length PQN-59::GFP exhibits slower diffusion rates that soluble GFP or GFP-PQN-59 fusion proteins lacking the three prion domains. These FRAP experiments were performed in developing oocytes. Graph indicates the recovery rates for the three proteins. PQN-59::GFP also exhibited slow diffusion rates in hypodermal cells (Figure S3). Quantification of the average recovery rates for each GFP protein depicted in I. Grey regions indicate that standard error of measurements (SEM) for 10-15 photobleaching events in separate animals.

To determine how PQN-59 activity contributes to normal animal development, we obtained a deletion allele of the *pqn-59* gene, *tm2960,* that removes 420nt of the *pqn-59* locus (Figure 3A). This lesion deletes the fifth exon of *pqn-59*; creating a premature stop codon upstream of the regions of the gene coding for the “prion-like” domains. This mutation suppresses *col-19::GFP* mis-expression phenotypes in *lin-4(ma161)* animals *pqn-59(tm2960); lin-4(ma161)* (60% adult n = 40) (Figure S1). Western blots from homozygous *pqn-59(tm2960)* animals suggest that this mutation is a null allele of *pqn-59* that generates no detectible PQN-59 protein and protein products of the predicted size for *PQN-59(tm2960)* are not seen in *pqn-59(0); cshIs38[PQN-59::GFP]* (Figure S1). Homozygous *pqn-59(tm2650)* animals that segregate from a balancer strain develop very slowly and exhibit a severe reduction in brood size (Figures 3D and E). The reduction in fecundity includes a reduced capacity to fertilize oocytes as well as a reduction in embryonic and early larval viability. Surprisingly, we did not observe any appreciable alterations in post-embryonic cell lineage in *pqn-59* mutants or precocious expression of adult-specific transcriptional reporters (Table I). The slow growth, sterility and larval lethality phenotypes of *pqn-59(0)* mutations can be rescued with a single copy PQN-59::GFP translational fusion targeted to chromosome II but not by related transgenes that encode PQN-59::GFP alleles that lack the UBA-like or “prion-like” domains suggesting that both of these domains are required for PQN-59 functions (Figure 3E). While animals harboring PQN-59(ΔUBA)::GFP or PQN-59(ΔPrD1-3)::GFP transgenes as the sole *pqn-59* gene failed to exhibit normal brood sizes, they exhibited no alterations in adult-specific alae formation (Table S1)

Proteomic analysis of whole worm lysates indicate that PQN-59 is a relatively abundant protein ranking in the top 5% of proteins expressed in embryos and larva (Figure 3F) [25]. In order to examine PQN-59 expression, we generated single-copy, C-terminal GFP-tagged version of *pqn-59* at the endogenous locus. Animals expressing this allele exhibited wild-type development and fecundity. Examination of PQN-59::GFP indicates that it is expressed throughout development and in all somatic and germline cells (Figure 3 G and H) (see below). At the subcellular level, PQN-59::GFP is localized exclusively in the cytoplasm with a marbled distribution. We also noted that, during post-embryonic development, there is a transient increase in PQN-59 expression in all somatic blast cells during and immediately after cell divisions and that PQN-59::GFP expression is maintained throughout adulthood.

### The “prion-like” domains of PQN-59 exhibit LLPS properties

Aside from the structured amino terminal UBA-like domain, a majority of the PQN-59 protein is predicted to contain intrinsically disordered regions (IDRs) and share features with IDRs of several proteins known to facilitate protein condensate formation (Figure 4A) [26–28]. Specifically, the carboxy-terminal ∼300 amino acids of PQN-59 are predicted to harbor three “prion-like” domains (PLDs) that are characterized by stretches of amino acids that are disproportionately enriched in glutamine (Q) and asparagine (N) residues when compared to most proteins in the *C. elegans* proteome (Figure 4A and S3) [19]. “Prion-like” domain containing proteins have also been demonstrated to exhibit natural “amyloid-like” properties which enable them to form fibril structures *in vivo* [29]. As a consequence, proteins harboring these IDR and or “PrD-like” domains exhibit a number of interesting biochemical properties including the ability to be co-precipitated by biotinylated-isoxazole (b-isox), which forms crystals in a temperature-dependent manner in aqueous solution and co-precipitates diverse proteins harboring low complexity domains (LCDs) from cell lysates [30, 31]. We tested if PQN-59 is precipitated by b-isox by using two approaches. First, we demonstrated that b-isox precipitates PQN-59 from whole worm lysates (Figure 4B). Because PQN-59 could be co-precipitated with other endogenous *C. elegans* proteins that are directly precipitated by the b-isox compound, we purified a fragment of PQN-59 that includes the “prion-like” domains as a GFP fusion protein and demonstrated that these domains are also precipitated by b-isox compound (Figure 4C). The b-isox does not precipitate soluble GFP [31] or proteins from whole-worm extracts that cross react with anti-PQN-59 antibodies or the minor *E. coli* proteins that co-purify with GFP-PQN-59(PrD1-3) indicating the specificity of b-isox for proteins harboring LCD and PrDs.

To determine if the “prion-like” domains of PQN-59 also exhibit LLPS properties, purified solutions of recombinant GFP-PQN-59(PrD1-3) were mixed with Ficoll or dextran sulfate. When these crowding reagents were added, the solution became opaque and turbulent with an increased A_600_ absorbance (Figure 4D and E). When these samples were examined with fluorescence and differential interference contrast (DIC) imaging, GFP-PQN-59(PrD1-3) solutions containing concentrating reagents exhibit micron-sized spherical droplets that freely move in solution and wet the surface of the glass coverslips (Figure 4F). Previous studies have used the aliphatic alcohol 1,6-hexandiol to probe the material properties of proteins that exhibit LLPS [32]. 1,6-hexandiol is thought to disrupt the weak hydrophobic protein-protein interactions that are required for the formation and stabilization of protein condensates [33, 34]. Incubation of GFP-PQN-59(PrD1-3) condensates with 1,6-hexandiol dramatically lowers the turbidity and absorbance of GFP-PQN-59(PrD1-3) solutions and eliminates visible condensate formation in microscopy samples (Figure 4D-F). To fully demonstrate that GFP-PQN-59(PrD1-3) condensates exhibit liquid-liquid-like dynamic properties we performed fluorescence recovery after photobleaching (FRAP) experiments. As demonstrated in Figure 4G, GFP-PQN-59(PrD1-3) condensates formed in 10% dextran sulfate solution rapidly recover fluorescence when a portion of the condensate is bleached (n>10). This rapid recovery was also seen in condensates formed with 150mg/mL Ficoll (n = 20) (Movie S1) indicating that GFP-PQN-59(PrD1-3) forms phase-separated liquid droplets *in vitro* (Figure 4G). Interestingly, prolonged incubation of these condensates (> 1 week) at 4°C leads to the formation of a hydrogel-like material that was incapable of being re-solubilized at room temperature in aqueous buffers (Figure 4H).

### Under normal growth conditions, PQN-59::GFP exhibits reduced mobility in the cytoplasm and this feature requires the “prion-like” domains

We next sought to examine the *in vivo* properties of PQN-59::GFP by comparing its diffusibility to that of a soluble, monomeric GFP. To accomplish this, we performed FRAP experiments using a strain harboring a single copy PQN-59::GFP (*cshIs38*) or a transgene driving soluble GFP driven from the *glh-1* promoter. We focused on measuring fluorescence recovery in developing oocytes where both proteins are localized in large and accessible cytoplasmic compartment (Figure 4I). As would be expected for a small, soluble protein, photobleached regions rapidly recover fluorescence signal in strains harboring soluble, monomeric GFP (Figure 4I). In contrast, photobleached cytoplasmic regions in oocytes expressing PQN-59::GFP recover fluorescence exceptionally slowly suggesting that PQN-59::GFP normally exhibits a limited diffusibility *in vivo*. We also performed FRAP on PQN-59::GFP in hypodermal cells and found that recovery was also slower than that observed for similar experiments with soluble GFP (Figure S2). We then sought to determine if the “prion-like” domains contributed to this feature. We integrated a PQN-59::GFP transgene that lacked the three PrDs, *cshIs78[pqn-59(ΔPrD1-3)::GFP*], at the same site where we targeted full length PQN-59::GFP above. This fluorescent reporter recovered faster than the full-length PQN-59::GFP reporter indicating that the PrDs of PQN-59 contribute to its reduced diffusibility in the cytoplasm (Figure 4I).

### PQN-59 exhibits LLPS *in vivo* and is required for efficient stress granule formation in developing oocytes

When cells are exposed to a variety of averse conditions, eukaryotic cells respond by dramatically reducing protein translation and re-distributing the localization of mRNAs and many mRNA-binding proteins [35]. Many stresses, including heat and chemicals, induce the formation of ribonucleoprotein complexes, called stress granules, that are formed from pools of untranslated mRNPs. These granules are dynamic, require a number of core proteins for formation and, when formed, show liquid-like behaviors [35]. The human orthologs of PQN-59, UBAP2 and UBAP2L, share a similar domain structure, are predicted to harbor “prion-like” and IDR domains (Figure S3) and have been demonstrated to localize to and function in the formation of stress granules in a variety of averse cellular conditions [22, 23, 36]. To determine if PQN-59 also functions in stress granule formation, we subjected animals expressing PQN-59::GFP to heat stress and examined PQN-59::GFP localization in developing oocytes. Heat stress (33°C) lead to the robust redistribution of PQN-59::GFP from its normal marbled, cytoplasmic localization to a large number of cytoplasmic puncta that vary in size from approximately 0.25-3.5µM (Figure 5A). To determine if the puncta observed in heat shocked oocytes were indeed stress granules, we also imaged the localization of the core stress granule component and sole G3BP1/2 ortholog in *C. elegans*, GTBP-1, that directly interacts with PQN-59 [37]. Under these conditions, PQN-59::GFP localization completely overlapped with the localization of GTBP-1::tagRED indicating that PQN-59 is a component of these structures during stress (Figure 5A). This same PQN-59::GFP transgene was localized to similar granules by a variety of stress treatments including arsenite treatment [37]. We used FRAP analysis to demonstrate that the PQN-59::GFP condensates that are induced by heat treatment exhibit LLPS properties as these cytoplasmic structures regain fluorescence rapidly after bleaching. As demonstrated in Figure 5B, photobleached PQN-59::GFP granules rapidly recovered fluorescence at a rate that is much faster than the recovery of bleached regions of PQN-59::GFP in normal growth conditions.

**Figure 5.**
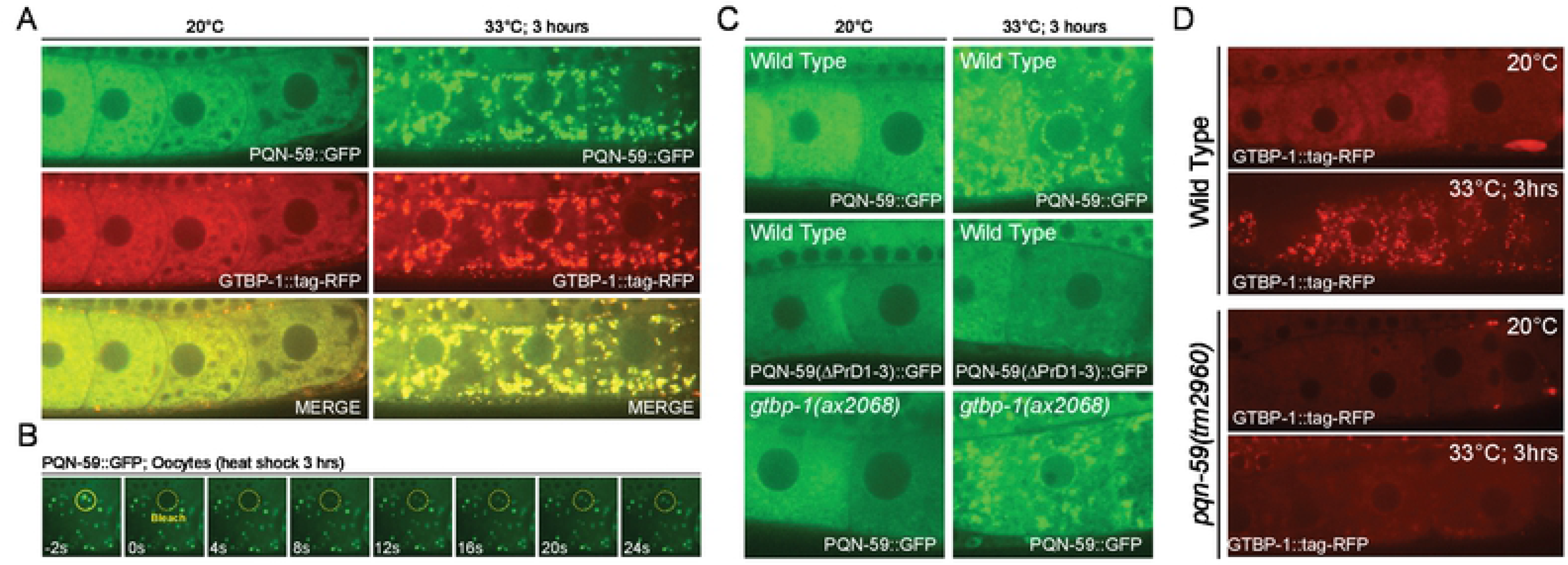
The “prion-like” domains of PQN-59 are essential for LLPS properties during heat stress and PQN-59 is important for the efficient localization of GTBP-1 to stress granules. **(A)** In developing oocytes, PQN-59::GFP is localized to the cytoplasm. At elevated temperatures, PQN-59::GFP re-distributes to cytoplasmically localized stress granules that co-localize with the core stress granule component, GTBP-1::RFP. **(B)** FRAP analysis of heat stress-induced PQN-59::GFP indicate that it is likely in a liquid-liquid phase separated state as PQN-59::GFP droplets recover fluorescence very rapidly when compared to PQN-59::GFP recovery at normal physiological temperatures (Figure 5I). **(C)** Condensation of PQN-59::GFP in heat stress conditions does not require GTBP-1 expression but does require the C-terminal “prion-like” domains of PQN-59. **(D)** GTBP-1 localization to stress granules in heat-shocked oocytes is severely compromised in the absence of PQN-59.

We next sought to determine what are the genetic and structural requirements of PQN-59 and GTBP-1 that are required for the localization of these proteins to heat-induced stress granules. First, we compared the ability of wild-type PQN-59::GFP to localize to stress granules to the dynamics of a version of PQN-59::GFP that lacked the “prion-like” domains. These experiments revealed that PQN-59(ΔPrD1-3)::GFP failed to localize to stress granules suggesting that the prion-like domains we had previously characterized as exhibiting LLPS properties *in vitro* are required for the LLPS properties of the full-length protein *in vivo* during stress granules formation (Figure 5C). In other systems, G3BP proteins are important for the assembly and stabilization of stress granules during heat stress [36, 38]. It is thought that G3BP1 in human cells is a central node of the RNA-protein network that triggers the formation of stress granules and this property is directly regulated by RNA binding [39, 40]. We tested whether PQN-59::GFP could be localized to stress granules in the absence of GTBP-1 expression by performing heat shock experiments in *gtbp-1(0)* animals. Animals lacking *gtbp-1* express normal levels of PQN-59::GFP and PQN-59::GFP localizes to stress granules upon heat treatment (Figure 5C). Since a significant portion of PQN-59::GFP is localized to stress granules in the developing oocytes of *gtbp-1(ax2069)* animals, we then asked if GTBP-1::tagRED association with stress granules requires PQN-59 activity. These experiments revealed that GTBP-1::tagRED localization to stress granules is dramatically reduced in *pqn-59(0)* animals (Figure 5D) indicating that PQN-59 is required for the efficient localization of GTBP-1::tagRED to these membranless organelles. Given the close association between PQN-59 and GTBP-1 and the importance of these proteins in stress granule formation, we tested whether *gtbp-1* (or other stress granule components) may also function in regulating temporal patterning during larval development. First we monitored adult specific alae formation in *gtbp-1(0)* mutants and found these cuticular structures were expressed as in wild-type animals (n = 32). We further tested whether the *gtbp-1(0)* mutation could suppress the reiterative heterochronic phenotypes in *lin-4(ma161)* mutants. Compared to the substantial suppression by *pqn-59(tm2960)*, combining *gtbp-1(0)* mutations with *lin-4(ma161)* had almost no detectible suppression of any alae phenotypes (adult-specific alae: *lin-4(ma161)* = 0 % alae (n = 40); *lin-4(ma161); gtbp-1(ax2068)* = 5% alae (n = 38); *pqn-59(tm2960); lin-4(ma161)* (45% adult n = 40). Depletion of two other stress granule components, *tiar-1 and tiar-2* [41–43], by RNAi (n > 30 each) also failed to suppress heterochronic phenotypes in *lin-4(ma161)* mutants suggesting that proper stress granule formation is distinct from roles in controlling temporal patterning.

### PQN-59 and UBAP2L interact with similar cellular components that are associated with stress granule formation, translation and post-transcriptional gene regulation

To gain insight in to the developmental function of PQN-59 in gene regulation, we sought to identify additional proteins that PQN-59 may function with to control developmental gene expression. To accomplish this, we immunoprecipitated PQN-59 from embryos using anti-PQN-59 antibodies and identified associated proteins via mass spectroscopy. Triplicate experiments identified 304 proteins that were reproducibly precipitated with PQN-59 antisera (Q value <0.5) (Table S2). We then performed gene ontology (GO) term enrichment analysis of PQN-59 co-precipitated proteins [44]. This analysis indicates that PQN-59 complexes with diverse sets of protein complexes including those enriched in RNA binding (mRNA and rRNA), protein translation (ribosomal proteins, translational initiation, elongation and regulatory factors), transcription (Direct DNA binding factors and co-activators), and transport (nuclear pore complex, motor proteins, microtubule binding) (Figure 6A). In addition to core complexes involved in translation, PQN-59 interacts with ALG-1, a core miRISC component whose mutation elicits phenotypes that are suppressed by *pqn-59(RNAi)* (Figure 2), and GTBP-1 which colocalizes with PQN-59 in stress granules (Figure 5).

**Figure 6.**
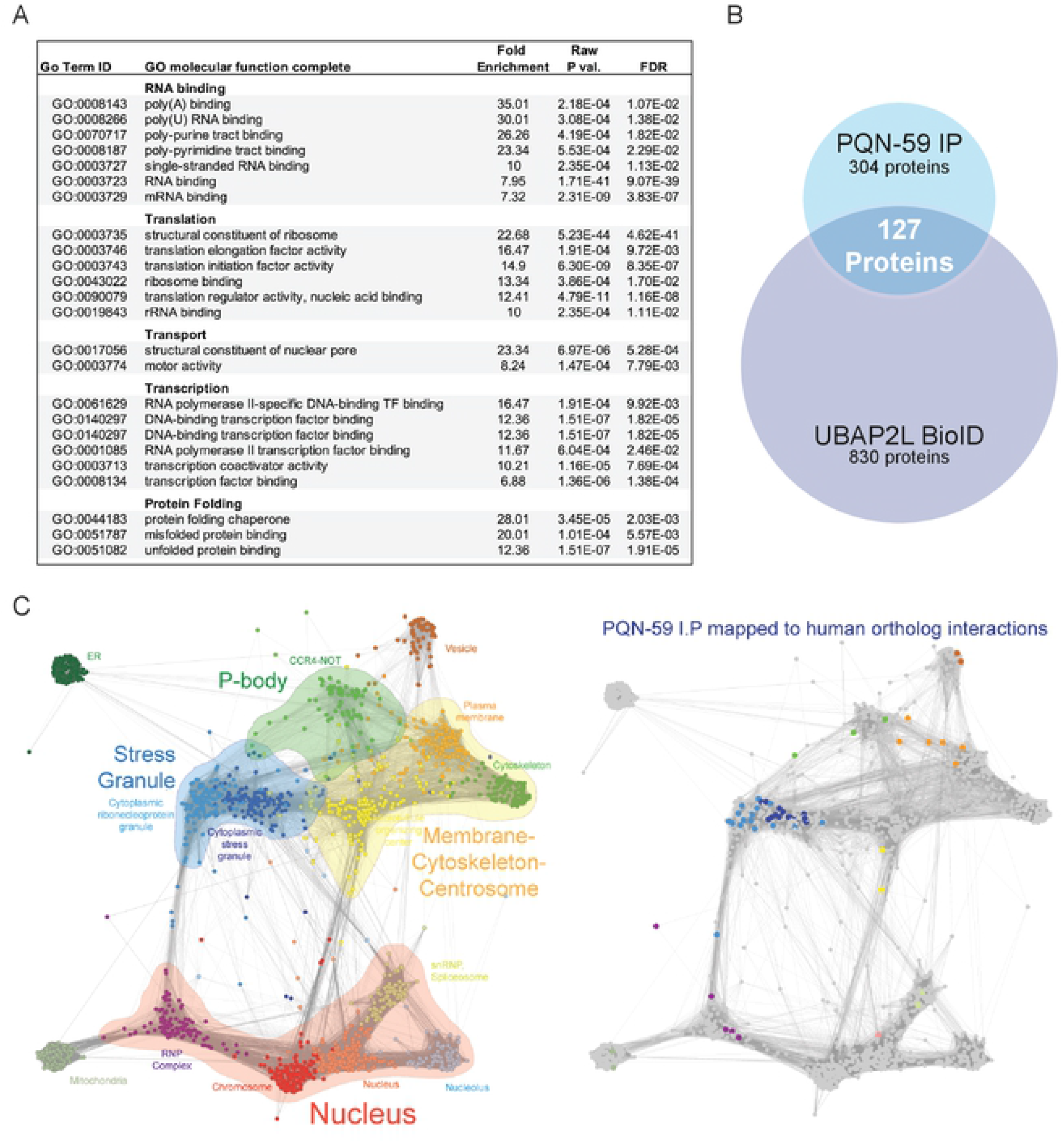
PQN-59 interacts with a variety of proteins involved in a variety of molecular processes and many of these interactions are shared with UBAP2L, the human ortholog of PQN-59. **(A)** Molecular function GO terms enrichment of PQN-59 interacting proteins isolated from developing embryos. **(B)** A Venn diagram indicating the overlap of orthologous proteins identified in PQN-59 I.P.s and BioID experiments performed with UBAP2L in human cells outlined in Youn et al. [24]. (C) The non-negative matrix factorization (NMF) network of protein complexes identified through massively-parallel BioID assays of RNA binding proteins derived from Youn et al. (left) [24]. Each group is organized by representative GO cellular compartment term as previously described [24]. The right network illustrates the location of PQN-59-interacting proteins that have orthologs that map to the indicated compartments.

The structural relationships between PQN-59 and human UBAP2 and UBAP2L (and Lingerer) (Figure S3) as well as their relationship to stress granule formation suggests that these proteins may function in an orthologous manner to regulate gene expression and the formation of stress induced RNA condensates. To determine if PQN-59 and UBAP2L interact with similar types of cellular components, we compared our list of PQN-59 interacting proteins to the list of proteins that were found to physically interact with UBAP2L as measured by BioID assays [24]. This molecular approach utilizes a promiscuous biotin ligase fused to a protein of interest to covalently tag proximally associated proteins *in vivo* [45]. When this approach was used to identify proteins that function near UBAP2L *in vivo*, a total of 830 interacting proteins were identified above background [24]. To compare these two lists, we employed the Ortholist server to identify orthologous pairs of human and *C. elegans* proteins from these lists (Table S3 and S4) [46, 47]. In addition to extensive, shared interactions between PQN-59 and UBAP2L with ribosomal subunits (Large Ribosomal subunits 2-7, 9-17, 18-22, 24-26, 28, 30, 33 and Small Ribosomal Subunits 0-3, 7-13, 17, 20, 23, 24, 27, 30), these efforts identified 127 orthologous pairs of proteins (Figure 6B); indicating that PQN-59 and UBAP2L are associated with overlapping complexes in vivo. We also noted that a large number of PQN-59 interacting proteins that do not share orthologous interactions with UBAP2L are enriched in GO terms associated with germline- and early embryonic-specific functions; proteins that would normally perform tissues specific functions not employed in human cell lines that are derived from somatic sources (Figure S4).

In order to both visualize PQN-59 interacting proteins into functional groups, we took further advantage of the large-scale application of the BioID proteomic approach outlined by Youn et al. that enabled an ultrastructural view of protein-protein interactions to be organized according to functionally and spatially related RNP-associated complexes (Figure 6C) [24]. When we overlayed the 127 PQN-59 interacting proteins that share orthologous protein-protein interactions with UBAP2 onto this map, we found that that a majority of proteins that are co-precipitated with PQN-59 are associated with other proteins that are found in two types of cytoplasmic condensates: stress granules and P-bodies (Figure 6C). While micron-sized stress granules are typically only visible during averse cellular conditions, multiple lines of evidence indicate that the protein-protein interactions that compose these bodies pre-exists in smaller forms during non-stress conditions [23, 24]. Furthermore, core miRISC components are functionally associated with both p-bodies and stress granules where they are thought to function in post-transcriptional regulation of miRNA target mRNAs [48].

### Depletion of *pqn-59* via RNAi alters the steady state levels of several miRNAs involved in temporal patterning

One mechanism by which *pqn-59* depletion could suppress the loss-of-function phenotypes associated with reduced miRNA activity would be to alter miRNA metabolism in a manner whereby the expression levels of these regulatory molecules are increased when PQN-59 expression is reduced. To test this hypothesis, we measured the levels of multiple miRNAs in animals that were exposed to control or *pqn-59* dsRNAs and extracted total RNA from late L4-staged animals. We then examined both miRNA processing and steady state levels using quantitative northern blots. As shown in Figure 7A, RNAi-mediated depletion of *pqn-59* increases the relative abundance of multiple, fully-processed miRNAs including those that regulate temporal patterning (*lin-4* and *let-7*). In addition, *pqn-59* depletion altered the levels of other mature miRNA species that are expressed in distinct tissues and temporal expression patterns [49]. Increases in mature miRNA levels during *C. elegans* development have been demonstrated to arise by elevated expression of miRNAs at the transcriptional level [6, 50] or by dampening mature miRNA turnover [51, 52]. If *pqn-59(RNAi)* alters mature miRNA levels by increasing the transcription of affected miRNA genes without improving pre-miRNA processing rates, we would expect pre-miRNA levels to increase in the absence of PQN-59 expression. A comparison of the pre-miRNA levels in control and *pqn-59(RNAi)* conditions indicate that pre-miRNA levels are not generally increased in the absence of PQN-59 (Figure 7A). Because pre-miRNAs for many miRNAs are efficiently processed in wild-type animals (precluding their relative accumulation), we also measured how depleting *pqn-59* expression alters pre-miRNA and miRNA levels in animals that harbor null mutations in *alg-1*, encoding one of the two *C. elegans* miRNA-specific argonaute proteins required for normal processing and stabilization of mature miRNAs [53]. Animals lacking ALG-1 expression, *alg-1(gk214)*, exhibit mild heterochronic phenotypes and accumulate the precursor mRNAs for a number of miRNA genes (Figure 7A) [53, 54]. Depletion of *pqn-59* in *alg-1(gk214)* animals suppresses the previously described and relatively weak heterochronic phenotypes (30% gapped alae in control RNAi conditions and 0% in animals exposed to *pqn-59* dsRNAs). As demonstrated in Figure 7A, depletion of *pqn-59* in *alg-1(gk214)* animals leads to an increase in mature miRNA species for a number of miRNAs without altering the levels of pre-miRNAs. Changes in mature miRNA levels in *alg-1(gk214); pqn-59(RNAi)* animals were validated using TaqMan assays of independent RNA samples (Figure S5A). Changes in mature miRNA levels for these miRNAs are not caused by an increase in the levels of two of the main miRISC components, ALG-1 and AIN-1, as levels of these proteins are not altered by PQN-59 depletion (Figure S5)B. These combined results suggest that mature miRNAs are stabilized in the absence of PQN-59 and this increase may enable specific miRNA targets to be more efficiently regulated in these conditions.

**Figure 7.**
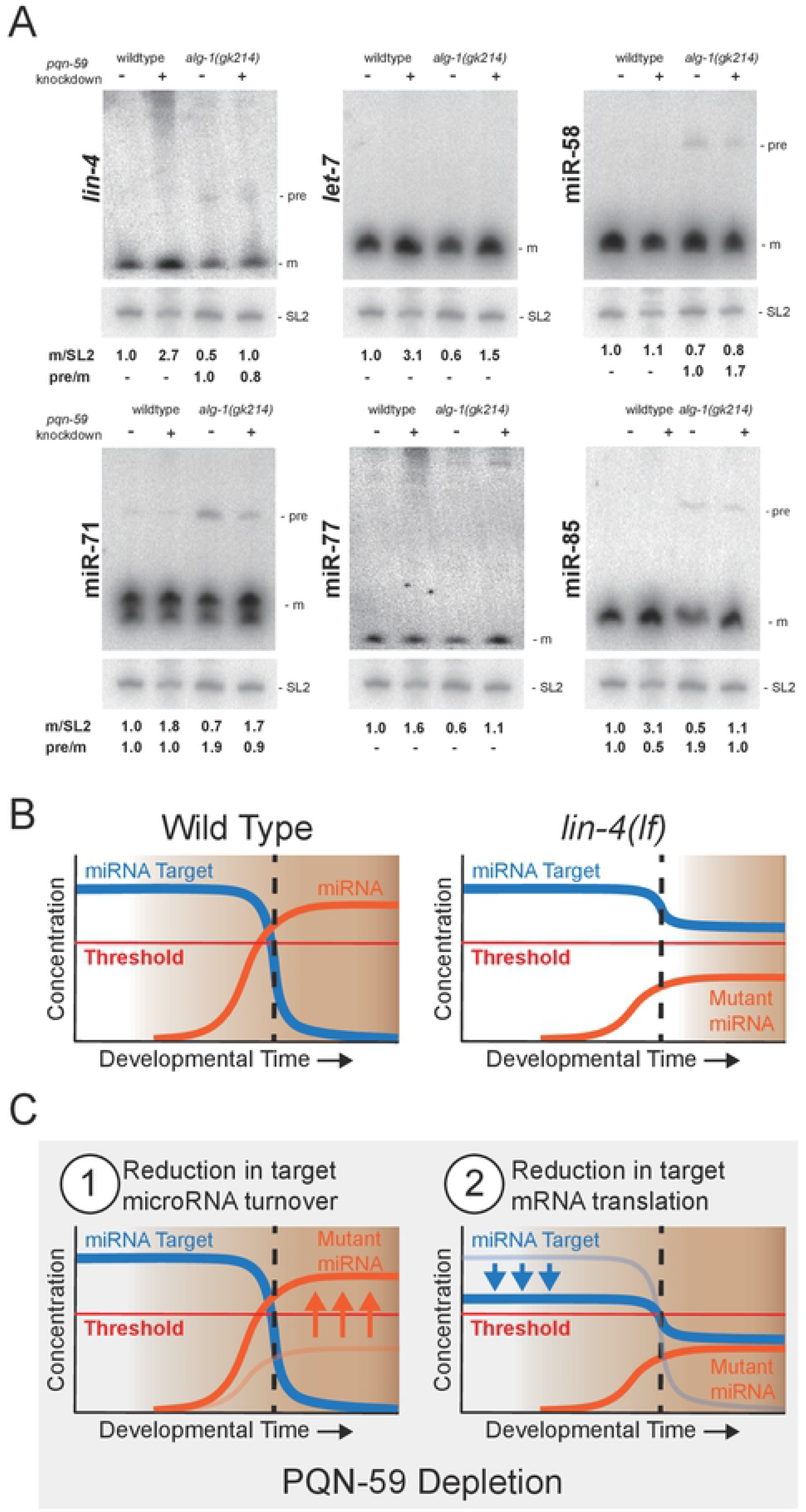
Depletion of *pqn-59* by RNAi increases the abundance of mature miRNAs. **(A)** miRNA northern analysis of total RNA extracted from staged, wild-type and *alg-1(0), alg-1(gk214),* mutant animals. Animals were exposed to control or *pqn-59* dsRNAs from hatching to the late L4 stage of larval development prior to RNA extraction. For calculation of relative expression of mature miRNA species, each experimental condition was calibrated to the expression level of 20µg of total RNA derived from wild-type animals exposed to control dsRNA. SL2 RNA levels were used as an input control. Pre-miRNA to mature RNA was calculated relative to the ratio of the signal in indicated sample. (B) A model of the generic bi-stable switch that incorporates a miRNA to control temporal cell fates during development. Normally (left side), the miRNA target gene is expressed above a critical threshold early in development and is responsible for enforcing earlier cell fate identities. At a defined point in development, the induced expression and activity of a miRNA gene rapidly curtails the target expression below a critical threshold enabling a clear change in cell fate at an important developmental milestone (demarcated by vertical dashed line). In miRNA mutants that under-accumulate functional miRNAs (e.g., *lin-4(ma161)*), the expression and activity of a mutant miRNA lowers the expression of the target gene but not below the critical threshold to enable a change in the bistable switch. (C) We propose two non-exclusively mutual mechanisms by which depletion of pqn-59 may suppress the defects associated with ineffective miRNA-mediated gene regulation in bistable developmental switches. In Model 1, depletion of *pqn-59* may increase the stability of the mutant miRNAs enabling the target gene to be efficiently dampened below the threshold at the developmental milestone. This model is consistent with data shown in above in panel A. Alternatively (Model 2), PQN-59 may function like UBAP2L and stimulate translation of some miRNA target mRNAs. Depletion of *pqn-59* would lower the expression of the target gene independent of miRNA expression. This reduced expression would nearer to the critical threshold needed to initiate a bistable switch in cell fate. In this scenario, the residual activity of the mutant miRNA may now be enough to reduce targe mRNA expression further and now below the critical threshold; enabling cell fate transformation in a bistable fashion.

## DISCUSSION

Bistability is a recurrent motif in developmental biology whereby distinct cell fates, defined by coherent patterns of underlying gene expression, can be switched by the activity of a key regulatory molecule within an established GRN. A fundamental property of bistable gene regulatory networks centers around changes in the temporal expression levels of specific, regulatory factors and the control of these levels around a critical threshold. Above this critical threshold, one cell fate is stable and below it, a distinct cell fate or expression program is enforced. In this manuscript we characterize the *C. elegans pqn-59* gene, encoding a conserved RNA-associated protein, as a component that functions to assure that cell fate specification events mediated by miRNAs during post-embryonic development exhibit bistability. We identified *pqn-59* in a reverse genetic screen aimed to identify suppressors of the severe temporal patterning phenotypes associated with hypomorphic alleles of two heterochronic miRNAs and a unique allele of an essential miRISC component ALG-1. While *pqn-59* depletion via RNAi can efficiently suppress these unique alleles, we also provide genetic evidence that *pqn-59* is not a bypass suppressor of *lin-4* and *let-7* (i.e., depletion offers little or no suppression of phenotypes associated with null alleles of these genes); indicating that PQN-59 normally functions as a modulator of miRNA activity. Consistent with this interpretation, treatment of wild-type animals with dsRNAs against *pqn-59* does not elicit detectible heterochronic phenotypes.

In order to illuminate mechanisms by which PQN-59 may function to modulate heterochronic gene expression, we analyzed phenotypes associated with a deletion allele of *pqn-59* gene and also characterized the expression and molecular features of PQN-59 *in vivo* and *in vitro*. Several lines of evidence suggest that *pqn-59* activities are not limited to regulating features of postembryonic temporal patterning and likely reflect a general role for this protein in development. First, deletion of the *pqn-59* gene results in pleiotropic phenotypes including an increase in the length of larval development and a dramatic reduction in fecundity. Furthermore, in contrast to the dynamic expression patterns of other heterochronic genes, PQN-59 is a very abundant and ubiquitously expressed protein. Third, depletion of *pqn-59* leads to the relative stabilization of several mature miRNAs species and does so without dramatically altering the levels of ∼70-nt pre-miRNA precursors. Because miRNAs are predicted to regulate the expression of a large fraction of expressed mRNAs [55, 56], inactivation of *pqn-59* may disrupt large portions of gene expression of which some, for example the heterochronic pathway, are exquisitely sensitive to perturbation.

The human ortholog of PQN-59, UBAP2L, is an established RNA-binding protein implicated in multiple biological processes. Large scale proteomic analysis of UBAP2L-associated proteins indicates that UBAP2L is highly integrated with other RNA-binding protein complexes implicated in stress granule formation and p-bodies, two membraneless cytoplasmic organelles formed through LLPS of RNAs and a multitude of RNA-binding proteins [23, 24]. UBAP2L expression is required for the formation of stress granules in adverse conditions [22–24]. Both p-bodies and stress granules are highly integrated with RNA-binding proteins that play direct roles in controlling gene expression and RNA turnover; including proteins directly involved in miRNA activity (e.g., miRISC components) [48]. Furthermore, UBAP2L appears to be directly associated with translating ribosomes and can be crosslinked to ribosomal RNAs and the coding sequences of hundreds of mRNAs; indicating that it may play a direct role in controlling translation [21]. Consistent with this hypothesis, artificially tethering UBAP2L to target mRNAs stimulates translation and depletion of UBAP2L results in a decrease in overall translational rates and a reduction in the levels of polysomes [21].

While the level of sequence identity between PQN-59 and UBAP2L is limited (18.6% Identity, 28.0% Similarity (EBLOSUM62)) (Figure S3), several lines of evidence suggest that these proteins perform orthologous functions. First, PQN-59, like UBAP2L, is predicted to lack overall strong secondary structure and is precipitated by biotinylated-isoxazole, a characteristic of proteins that harbor exhibit LLPS features in vivo [31]. We also demonstrate that the prion-like domains of PQN-59 themselves exhibit LLPS properties *in vitro* and that during heat stress, PQN-59 localizes to stress granules and exhibits LLPS properties *in vivo*. Structure-function analysis of PQN-59 domains indicate that the prion-like sequences in the C-terminus of PQN-59 are required for stress granule localization. Second, like UBAP2L, PQN-59 expression is required for the efficient recruitment of GTBP-1, the sole *C. elegans* G3BP ortholog, to stress granules [23, 24, 36]. Third, condensation of PQN-59 into stress granules is not dependent on GTBP-1; further mirroring the epistatic relationship between these proteins in human cells [22]. Finally, biochemical characterization of PQN-59 associated proteins indicate that UBAP2L and PQN-59 are physically associated with similar protein complexes in vivo; suggesting that they both may integrate aspects of mRNA metabolism, regulation and expression through these interactions.

Our genetic characterization of *pqn-59* during *C. elegans* development and the potential conservation of functions between PQN-59 and UBAP2L enable us to propose at least two non-mutually exclusive models for the function of PQN-59 in temporal patterning. A cornerstone of miRNA function in the heterochronic pathway centers on the rapid downregulation of their target mRNAs below a critical threshold (Figure 7B). In contrast to what happens in wild-type animals, animals lacking full activity of heterochronic miRNAs (as exemplified for *lin-4(ma161)* mutants) are unable to dampen the expression of their target mRNA below critical threshold required for the bistable transition in temporal cell fate (Figure 7B). As a consequence, *lin-4(ma161)* animals exhibit heterochronic phenotypes that are indistinguishable from those in animals that completely lack *lin-4* (as exemplified for *lin-4(e912)* mutants) [6]. As demonstrated in Figure 7A, *pqn-59* depletion results in the stabilization of many mature miRNAs. We hypothesize that the potentially generalized stabilization of mature miRNAs elicited by depleting *pqn-59* expression may enable the levels of critical miRNAs to increase to a level where they can now effectively dampen *lin-14* expression (Figure 7C model 1). In addition to this potential mechanism, PQN-59, like UBAP2L, may promote general protein translation. In this context, depletion of *pqn-59* reduces the normal expression levels of miRNA target genes (like *lin-14*) to a level that is closer to the threshold that defines the bistable switch between cell fate specification (Figure 7C model 2). Therefore, *pqn-59* may function to normally assure that this bistable switch that defines the L1 to L2 transition in *C. elegans* larval development is not inappropriately crossed unless a sufficient miRNA is expressed. Both of these models are consistent with *pqn-59* functioning outside of the normal heterochronic GRN and furthermore explain the observation that *pqn-59(RNAi)* is an efficacious suppressor of multiple miRNA loss-of-function phenotypes but that *pqn-59* depletion is incapable of bypassing the activities of these genes.

Overall, our study demonstrates that *pqn-59* functions to modulate gene expression and cell fate specification during *C. elegans* development. Prior studies have indicated that UBAP2L and its functions with other stress granule and RNA-binding protein partners may play a complex role in a variety of human diseases. These include a role for UBAP2L overexpression in various types of cancer [57–62] and in pathologies that are related to protein aggregation in neurodegeneration [63–66]. We imagine that the genetic and experimental tractability of the *C. elegans* model will be instrumental in discovering the underlying mechanisms by which this conserved family of proteins functions.

## SUPPLEMENTAL FIGURES, TABLES and MOVIE LEGENDS

**Figure S1. *pqn-59(tm2960)* phenocopies *pqn-59* RNAi in suppressing *lin-4(ma161)* heterochronic phenotypes and is likely a null allele. (A)** *pqn-59(tm2960); lin-4(ma161)* animals exhibit wild type *col-19::GFP* expression (and protruding vulva phenotype) while animals harboring a single copy of the *pqn-59* deletion allele (balanced with an *hT2 myo-2::GFP* balancer) exhibit only a very mild *col-19::GFP* expression phenotype and no vulval induction. (B) Western blots of wild-type, cshIs25[PQN-59::GFP CRISPR allele at pqn-59 locus], and *pqn-59(tm2960); cshIs38* [PQN-59::GFP single copy on Chromosome II] animals using antibodies against the PQN-59 amino terminus.

**Table S1. Quantification of heterochronic phenotypes of homozygous *pqn-59(tm2960)* animals expressing variations of PQN-59::GFP deletion alleles targeted to chromosome II.** Assays include measurement of L4 stage and adult alae formation.

**Figure S2. FRAP of PQN-59::GFP in hypodermal cells indicate that PQN-59 exhibits a reduced diffusion rate compared to soluble GFP. (A)** FRAP analysis of soluble GFP (driven from the dcap-1 promoter) and a translational fusion of PQN-59. **(B)** Quantification of the recovery rates for each GFP protein depicted in A. Graphs represent the average recovery rate and error bars indicate that standard error of measurements (SEM) for 10-15 photobleaching events in separate animals.

**Movie S1. Liquid-Liquid Phase Separated droplets of GFP-PQN-59(PrD1-3) exhibit intra-droplet flow when photobleached.** One half of two recently fused GFP-PQN-59(PrD1-3) droplets were subjected to FRAP and monitored for fluorescent recovery.

**Figure S3. Comparison of the “prion-like” domain structures and predicted folding characteristics of the *Drosophila* and human PQN-59 orthologs.** The PLAAC: Prion-Like Amino Acid Comparison Server (http://plaac.wi.mit.edu) was used to query the primary amino acid sequences of PQN-59, Lingerer isoform A, UBAP2 and UBAP2L with the following parameters: Core length of 60 and a relative weighting of background probability set to the corresponding species of origin. In protein sequences below each PLAAC graph, red highlighted amino acids indicate glutamine (Q) and asparagine (N) rich sequences identified by this analysis as encoding “prion-like” domains.

**Table S2. Summary Table outlining various PQN-59 interacting proteins measured by Mass Spectroscopy.** This table categorizes PQN-59 interacting proteins (peptides), gene name, and molecular descriptions. This table also quantifies the enrichment, percent change, P values, and Q values from these triplicate PQN-59 I.P. experiments from embryo extracts.

**Table S3. List of orthologous human gene names for proteins immunoprecipitated in PQN-59 complexes outlined in Table S1.**

**Table S4. List of orthologous C. elegans gene names of proteins identified as UBAP2L interacting proteins in BioID experiments outlined in Youn et al. 2018.**

**Figure S4. PQN-59 Interacting proteins that do not map to UBAP2L-BioID interactors are enriched for GO terms associated with germline and fertility.** Relative GO term enrichment (molecular function) scores for proteins that co-precipitate with PQN-59 from early embryos.

**Figure S5. *pqn-59* depletion corrects the reduction of mature *lin-4* and *let-7* miRNA levels observed in *alg-1(0)* mutants and has little effect ALG-1 or AIN-1 expression (encoding two of the major miRISC components). A)** Taqman analysis of *lin-4* and *let-7* miRNAs isolated from wild-type or *alg-1(gk215)* animals subjected to control or *pqn-59* dsRNAs. In each measurement was standarized by also quantifying the expression of U18 snoRNA in each sample. Error bars represent standard deviation (n=3 biological replicates, two technical replicates). P-values were calculated using a Student’s t-test, and corrected for multiple comparisons using a Bonferroni correction. **(B)** In the *eri-1(ok2683)* RNAi hypersensitive strain *pqn-59* depletion decreases PQN-59 to 12% of its abundance in control *eri-1(ok2683)* animals. In wild-type animals *pqn-59* depletion via RNAi decreases PQN-59 to less than half of its abundance in animals fed control RNAi. In contrast, ALG-1 and AIN-1 abundance are minimally affected by *pqn-59* depletion. Western blots were prepared with LiCOR reagents and imaged with a Classic Infrared Odyssey imager. Quantification was performed in ImageJ.

**Table S5. List of *C. elegans* strains used in this study.**

## MATERIALS AND METHODS

### *C. elegans* maintenance and genetics

*C. elegans* strains were maintained on standard media at 20°C and fed *E. coli* OP50 [68]. Some strains were provided by the CGC, which is funded by NIH Office of Research Infrastructure Programs (P40 OD010440). *pqn-59(tm2960)* was obtained from Shohei Mitani the National BioResource Project (NBRP) at the Tokyo Women’s Medical University. Hypochlorite treatment followed by overnight starvation was used to synchronize animals at L1. *C. elegans* strains used in this study are listed in Table S5.

### RNAi feeding

RNAi by feeding was performed using *E. coli* (strain HT115) expressing double stranded RNA corresponding to endogenous *C. elegans* ORFs (SourceBioScience, United Kingdom) using standard methods [69]. This library included plasmids producing dsRNA against *pqn-59* or a control dsRNA expression plasmid (pPD129.36) [70] that does not contain sequence corresponding to any *C. elegans* gene [71, 72]. Positive scoring clones were verified by replicate experiments and dsRNA-targeted gene product was confirmed by Sanger sequencing. To prevent contamination by *E. coli* OP50, late L4, P_0_ animals were added to RNAi plates individually after removing co-transferred bacteria. Unless otherwise noted, F1 progeny were analyzed for RNAi-induced phenotypes following 3-4 days incubation at 20°C.

### Northern Blotting and miRNA Taqman Assays

Total RNA was isolated using TRIzol (Invitroen) from staged populations of worms that were exposed to control or *pqn-59* dsRNAs until the late L4 stage. Northern blots were performed as previously described [12, 73]. Taq man analysis of extracted RNAs and calculations were performed as previously described [74].

### CRISPR recombineering

The GFP-tagged allele of *pqn-59* at the endogenous locus (*cshIs25)* was created using standard CRISPR recombineering [75]. Homologous recombination was carried out with a modified version of the Cas9/Guide plasmid (pCMH1299) where the *pqn-59*-specific guide sequence, GTACAACTGGAGTAACTAAC, was inserted upstream of the single RNA guide backbone. Homologous repair and insertion of the GFP ORF used pCMH1304 that contained the GFP ORF plus approximately 1200bp 5’ and 1000bp 3’ flanking regions (centered on the endogenous *pqn-59* stop codon). Full-length(*csh38)* and deletion alleles (ΔUBA(*cshIs86)* and ΔPrD1-3(*cshIs78)*) of PQN-59::GFP were targeted to Chromosome II using repair templates that target the ttTi5606 site using pCMH1370, pCMH1375 and pCMH1603, respectively. CRISPR editing at the ttTi5606 target site was accomplished using guide sequences in pDD122 following standard approaches [75].

### Wester blotting and antibody production

Antibodies Against PQN-59 were made by immunizing separate rabbits with OVA or KLB conjugated peptides against amino acids 5-19 (GDKKATSDQARLARL) or 628-643 (PNLSSLFMQQYSPAPH) of the predicted PQN-59 protein. Western blots used the N-terminal antibody (targeting amino acids 5-19 (GDKKATSDQARLARL)). For the antibody used in IP experiments, a PQN-59 a C-terminal fragment (amino acids 304-712) was cloned using the Gateway technology (Invitrogen) into the pDEST15 and purified as previously described [37]. Whole worm lysates were prepared by dounce homogenizing staged, wild-type or transgenic animals in an equal volume of Pedro’s buffer (30mM HEPES, pH 7.5, 100mM KOac, 10mM EDTA, 10% glycerol, 1 mM DTT, 1x Roche complete mini protease inhibitors, Sigma phosphatase inhibitor cocktails 1 and 2 (each 1:100) and 1% (v/v) SuperRNase-IN). Lysates were clarified by centrifugation at 13,817 x g for 20 minutes at 4°C. Protein concentrations were measured using Bradford Assay (ThermoFisher, cat# 23200). Antibodies used in this study are as follows: anti-tubulin (Abcam, cat# EPR13478(B)), Anti-ALG-1 [15], anti-AIN-1 [76], TrueBlot HRP-conjugated anti-rabbit, anti-mouse and anti-Rat secondary antibodies (eBioscience).

### Recombinant protein purification

pCMH1726, encoding the pHIS6-PQN-59(PrD1-3)_YFP protein, was constructed using GIBSON cloning with a PCR fragment of the *pqn-59* cDNA (encoding the terminal 305 aminos acids of PQN-59) and pHIS6_Pararallele_GFP vector described in Kato et al.; replacing the Fus(LC domain) ORF in pHIS6-Parallel-FUS(LC)[77]. The His6::GFP::PQN-59(PrD1-3) protein induction and purification was accomplished using previously described protocols[77]. Briefly, BL21(DE3) + pRIPL cells were transformed with pCMH1726 (pHIS6-parallele-GFP::PQN-59(PrD1-3) and selected on LB + ampicillin + chloramphenicol plates. A single colony was grown overnight in selective media and re-inoculated into 1L of LB medium and grown to an O.D.600 of 0.6. The culture was then cooled on ice for 20 minutes and 0.5mL of 1M IPTG was added. Cultures were then grown overnight at 16°C. Bacteria were collected by centrifugation and lysed with a sonicator in 35mL of Lysis buffer (50 mM Tris-HCl pH 7.5; 500 mM NaCl; 1% Triton X-100; 20 mM β-mercaptoethanol (BME); 1 tablet of protease inhibitors (Sigma S8830, 1 tablet per 50 mL)). Samples were centrifuged at 35,000 rpm for 30 minutes at 4°C. The supernatant was removed to a fresh 50 mL conical and further incubated with 2mL of Talon Beads (Qiagen) for 1 hour at 4°C. Extract slurry was then applied to a column. Beads were washed with approximately 75-100 mL of lysis buffer (above) supplemented with 20mM imidazole. Samples were eluted in 2mL fractions of elution buffer supplemented with 250mM imidazole and quantified for concentration and purity using SDS-PAGE.

### Microscopy and fluorescence recovery after photobleaching (FRAP) assays

For imaging of *C. elegans* larva in figures 1 and 2, mages were acquired with a Zeiss Axio Observer microscope equipped with Nomarski and fluorescence optics as well as a Hamamatsu Orca Flash 4.0 FL Plus camera. An LED lamp emitting at 470 nm was used for fluorophore excitation. For single images, animals were immobilized on 2% agarose pads supplemented with 100mM Levamisole (Sigma). Photobleaching of phase separated droplets or transgenic animals were carried out using a laser on a Nikon Ti-E microscope fitted with a Perkin-Elmer UltraVIEW VoX high speed spinning disk (Yokogawa® CSU-X1) laser confocal microscope with live cell imaging capability, time-lapse microscopy, photokinesis, multi-position image acquisition, 6 diode laser lines (405, 440, 488, 514, 561 and 640nm). Imaging was performed at room temperature using spinning-disc confocal microscopy system (UltraVIEW Vox; PerkinElmer) and a charged-coupled device camera (ORCA-R2; Hamamatsu Photonics) fitted to an inverted microscope (Ti Eclipse; Nikon) equipped with a motorized piezoelectric stage (Applied Scientific Instrumentation). Image acquisition and analysis was performed using Volocity version 6.5 (Quorum Technologies). For droplets, LLPS samples were spotted onto 22mm x 22mm coverslips which were rapidly placed onto the top surface of a slide. After droplets had wetted to the glass surface, individual regions were photobleached. *C. elegans* samples were prepared as above and subjected to photobleaching in a similar manner to LLPS droplets. Appropriate ROI outside and inside samples (adjacent to bleached regions) were taken as controls. The selected region of interest (ROI) was bleached with a 100 % laser power. All the measurements were performed at room temperature and at least in triplicates. For *C. elegans* animals FRAP, the bleaching of ROI (5-10 μm) was performed at a 100% laser power and 50 iterations, after acquiring 2-20 images before bleaching. The recovery was monitored for ∼100-300 s. The images were corrected for laser bleaching by selecting a fluorescent region outside the ROI. FRAP data analysis as previously described [78].

### Proteomic analysis of PQN-59 complexes

N2 worms were grown on OP50-seeded NGM plates, and embryos were harvested from gravid worms by bleaching (500 mM NaOH, 15% bleach). Embryos were resuspended in IP buffer (100 mM KCl, 50 mM Tris pH 7.5, 1 mM MgCl_2_, 1 mM DTT, 5% glycerol, 0.05% NP40, 1 mM EDTA, Protease Inhibitor Cocktail (Roche)) and frozen in liquid nitrogen. For protein extraction, the embryos were ground on dry ice using a mortar and pestle. The embryonic protein homogenate was thawed on ice and centrifuged at 14,000 rpm for 30 minutes. Equivalent amounts of PQN-59 serum or pre-serum, for control, were incubated with 10 μl of Protein G UltraLink Resin (Thermo Scientific) on ice for 1 hour and 30 minutes. After three washing steps of the beads with IP buffer, approximately 2 mg of embryonic protein homogenate was added and combined samples were incubated for 2 hours on ice. Beads were then washed three times with IP buffer and an additional three times with a “last wash” IP buffer (100 mM KCl, 50 mM Tris pH 7.5, 1 mM MgCl_2_, 1 mM DTT, 1 mM EDTA). PQN-59 complexes were eluted in 0.15% trifluoroacetic acid (TFA). Isolated samples were then frozen on dry ice and subjected to mass spectrometry analysis. Three IP samples from two separate protein homogenates were used in the analysis.

### Bioinformatic analysis

Prion domains of the *pqn-59* ORF were identified using PLAAC (http://plaac.wi.mit.edu). Additional protein domain/motif prediction tools were used to identify other, conserved domains of PQN-59 including PROSITE at ExPASy (https://prosite.expasy.org/), MOTIF (GenomeNet, Institute for Chemical Research, Kyoto University, Japan) (https://www.genome.jp/tools/motif/), InterPro (http://www.ebi.ac.uk/interpro/) and SMART (http://smart.embl-heidelberg.de) [79]. GO term analysis was performed using PANTHER Tools (http://www.pantherdb.org) [80, 81].

## AKNOWLDEGEMENTS

We would like to thank Tim Schedl, Steve McKinght, Popi Syntichaki, Dustin Updike, and Geraldine Seydoux for strains and reagents used in this study. We are also indebted to the CSHL Bioinformatics Resource and the Shared Microscopy Core Resource (which are funded, in part, by the CSHL Cancer Center Support Grant 5P30CA045508) for aiding in the analysis of proteomic data and microscopy, respectively. Cold Spring Harbor Laboratory, the Rita Allen Foundation, HIH NIGMS R01GM117406 supported C.M.H..

## AUTHOR CONTRIBUTIONS

**Conceptualization:** Christopher M. Hammell, Robin Weinmann, Natalia Stec, Francoise Schwager, Simona Abbatemarco, Jing Wang, Ouyang Huiwu, Colleen Carlston, Monica Gotta

**Data Curation:** Christopher M. Hammell, Robin Weinmann, Natalia Stec, Francoise Schwager, Simona Abbatemarco, Jing Wang, Ouyang Huiwu, Colleen Carlston, Monica Gotta

**Formal analysis:** Christopher M. Hammell

**Funding acquisition:** Christopher M. Hammell and Monica Gotta

**Methodology:** Christopher M. Hammell and Monica Gotta

**Project administration:** Christopher M. Hammell and Monica Gotta

**Writing - original draft:** Christopher M. Hammell

**Writing -review and editing:** Christopher M. Hammell, Jing Wang, Ouyang Huiwu, Simona Abbatemarco, and Monica Gotta

